# The Effects of Motor Modularity on Performance, Learning, and Generalizability in Upper-Extremity Reaching: a Computational Analysis

**DOI:** 10.1101/804096

**Authors:** Mazen Al Borno, Jennifer L. Hicks, Scott L. Delp

## Abstract

It has been hypothesized that the central nervous system simplifies the production of movement by limiting motor commands to a small set of modules known as muscle synergies. Recently, investigators have questioned whether a low-dimensional controller can produce the rich and flexible behaviors seen in everyday movements. To study this issue, we implemented muscle synergies in a biomechanically realistic model of the human upper extremity and performed computational experiments to determine whether synergies introduced task performance deficits, facilitated the learning of movements, and generalized to different movements. We derived sets of synergies from the muscle excitations our dynamic optimizations computed for a nominal task (reaching in a plane). Then we compared the performance and learning rates of a controller that activated all muscles independently to controllers that activated the synergies derived from the nominal reaching task. We found that a controller based on synergies had errors within 1 cm of a full-dimensional controller and achieved faster learning rates (as estimated from computational time to converge). The synergy-based controllers could also accomplish new tasks–such as reaching to targets on a higher or lower plane, and starting from alternate initial poses–with average errors similar to a full-dimensional controller.

## 1 Introduction

Every movement we make, from picking up a glass of water to opening a door, requires the control of a large number of muscles, which is complex due to nonlinearities and redundancies in the musculoskeletal system. It has been hypothesized that the central nervous system (CNS) uses modularity to simplify the control of our high-dimensional musculoskeletal system, limiting motor commands to a small set of modules known as muscle synergies [10]. A muscle synergy is a group of muscles that have a fixed ratio of excitations. Evidence from experiments with lower vertebrates, cats, and monkeys suggests that muscle synergies are encoded in the spinal circuitry [7, 30, 33]. Previous investigators have also shown that the muscle activity measured from electromyography (EMG) can be reconstructed accurately from a linear combination of a small number of muscle synergies for a variety of tasks, including walking [8, 40] and reaching [11, 38]. However, other studies have presented evidence that synergies may be a by-product of task and performance constraints [15, 16, 39]. A better understanding of the advantages and limitations of muscle synergies may help us evaluate their evolutionary role across different species [19] as well as their role in motor development and learning [27].

More evidence is needed to test the hypothesis that the CNS uses muscle synergies to simplify control, particularly for movements of the upper extremity. Several investigators have argued for the need to assess muscle synergies based on task achievement rather than on their ability to reconstruct EMG, because small errors in muscle excitations could cause large errors in task performance [3, 16, 28]. In simulation studies, Berniker et al. [5] and Kargo et al. [31] showed that in the movement of the frog hindlimb, a low-dimensional controller implemented with muscle synergies causes negligible performance degradation. In another simulation study, Meyer et al. [37] showed that muscle synergy controls result in more accurate walking kinematic and kinetic predictions compared to independent muscle control. In contrast, for the upper extremity, previous computational work indicates that muscle synergies can introduce substantial aiming errors and limitations in terms of endpoint stiffness and energy consumption during isometric tasks [16, 28]. However, this assessment was made with simple biomechanical models with little redundancy, where any dimensionality reduction is likely to introduce substantial errors. For instance, in de Rugy et al. [16], synergies introduce larger aiming errors in their simplified wrist model compared to their complex elbow model.

Evidence that modularity aids motor learning could provide additional support for the muscle synergy hypothesis. For example, recent experiments with rats that underwent spinal transections provides evidence that synergies may be formed early postnatally, and the authors suggest that these synergies could be evolutionarily advantageous by simplifying skill acquisition through adulthood [53]. Muscle synergies might speed the learning process by restricting the size of the action space (i.e., the space of possible muscle excitation patterns) [48]. On the other hand, it is possible that a large action space could facilitate skill acquisition as it would provide an abundance of ways to produce the desired behavior [35]. In Berger et al. [4], after a virtual tendon rearrangement surgery, subjects had difficulty adapting to perturbations that were incompatible with synergies identified pre-virtual-surgery, whereas they quickly adapted to perturbations compatible with the synergies. More information about the role of motor modularity in learning new or more complex tasks is needed.

An additional compelling case for the synergy hypothesis would be if a few synergies could generate a wide variety of behaviors. There is some evidence for the generalizability of synergies from experiments in the frog hindlimb during jumping, swimming, and walking movements [10]. Researchers have also demonstrated that the synergies computed from EMG measured in one task can reconstruct EMG measured for other tasks, including postural perturbations in cats [50] and humans [9, 51], and isometric force generation in humans [42]. Similarly, it has been shown that only six synergies are necessary to reconstruct EMG accurately for planar reaching movements with different directions and speeds [11], and through visuomotor adaptations [22]. These studies provide promising evidence for the generalizability of synergies, but as discussed above, even small errors in muscle excitations could significantly degrade task performance. In simulation studies, researchers have shown that synergies computed for one walking task facilitate the generation of walking dynamics at different speeds [37] and body weights [36], but it is now known if these results extend to the upper extremity and non-periodic tasks. Giszter suggests that a possible hybrid control architecture, where the CNS has access to both synergies and the control of individual muscles, could potentially avoid the limitations of a reduced control set, while maintaining its advantages [23].

Computational models can help reveal the interplay between synergies and task space errors, motor learning, and generalization, allowing us to quantitatively assess these debated relationships. While computational studies have begun to unravel these relationships, previous models have often been simplified due to the challenges of modeling complex musculoskeletal structures and synthesizing realistic movements. Detailed models of the upper extremity are available [44], along with computational models that synthesize kinematics instead of track experimental data. These computational models are being used to study cause-effect relationships between muscular deficits and movement abnormalities and to test hypotheses in motor neuroscience. [2, 18, 20].

We implemented muscle synergies in a biomechanically realistic upper extremity model [44] and used optimal control to synthesize reaching movements to a specified target. We focus on point-to-point movements because many voluntary upper extremity movements consist of moving from one pose to another, and being able to hold one or more joints in a static pose [46]. We performed computational experiments to determine whether a modular architecture based on muscle synergies introduces task performance deficits, as defined by a cost term that incorporates target accuracy and muscle effort. We examined the rate the optimization converged to a set performance threshold for models with varying numbers of synergies to assess whether synergies facilitate motor skill acquisition. We also tested whether synergies generated from one set of reaching tasks generalized to new tasks, such as reaching from alternate starting postures or while holding a weight. Finally, we implemented a hybrid controller that included both synergy and individual muscle control to determine whether this architecture improved generalizability to new tasks without sacrificing convergence rates. Overall, our aim was to assess how muscle synergies influence the ability to acquire and generate a repertoire of movements.

## 2 Methods

### 2.1 Modeling and Simulation

We used an upper extremity biomechanical model developed by Saul et al. [44] (available at https://simtk.org/projects/upexdyn). It consists of 47 Hill-type muscle-tendon actuators [54] with parameters and paths derived from experimental and anatomical studies. The skeletal model has three degrees-of-freedom for the shoulder, one degree-of-freedom for the elbow, and one degree-of-freedom for the forearm to allow for pronation and supination. We removed the wrist degree-of-freedom. The biomechanical modeling and simulation was performed with OpenSim 3.3 [17] using a semi-explicit Euler integrator with an accuracy of 1e–2.

### 2.2 Trajectory Optimization

We used a numerical optimal control method to synthesize motion [1]. We optimized for muscle excitations that minimized the following cost function:

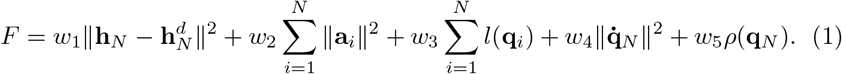

The terms **h**_*N*_ and 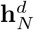 denote the actual and desired position of the center of the hand at the last timestep *N* of the trajectory, **a**_*i*_ denotes the vector of muscle activations at timestep *i, l*(**q**_*i*_) is a quadratic penalty on joint limit violations given pose 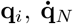 denotes the joint velocities at the last timestep, *ρ*(**q**_*N*_) is a quadratic term to encourage pronation of the forearm at the last timestep (as is typical in reaching movements to a target), and || · || denotes the 2-norm. The muscle activations are related to the muscle excitations through a first-order differential equation [54]. We empirically tuned the weights *ω*_1_, *ω*_2_, *ω*_3_, *ω*_4_ and *ω*_5_ to 5, 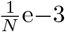, 1e-5, 0.15 and 0.5. We refer to 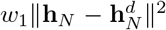 and 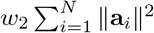 as the *task* and *effort* terms, respectively. The cost function encourages the hand to be as close as possible to a target with a small final velocity and the forearm in pronation, while minimizing muscle activations and joint limit violations throughout the movement. We specified the movement duration *T* and the number of timesteps *N* in the trajectory was chosen to achieve the desired integration accuracy.

We optimized for the muscle excitations to produce movement with the Covariance Matrix Adaption Evolution Strategy (CMA-ES) [26]. The free variables were the values of the muscle excitations at every 0.1 s interval in the movement. The excitations were held constant for the 0.1 s interval. The CMA-ES algorithm used a Gaussian distribution over the free variables to sample a number of possible control solutions (referred to as the population size). Given each sample of control solutions (i.e., muscle excitations), we used a forward simulation to obtain a trajectory which we evaluated according to the cost function described above. A new Gaussian distribution was then formed as a function of a subset of the samples with the lowest costs. The CMA-ES algorithm iterated through this process to steer the distribution towards a low cost region. The number of iterations and the population size in CMA-ES determined the exhaustiveness of the search; we set both to 100 based on empirical testing. We tested a more exhaustive search by doubling the population size and the number of iterations to ensure that it yielded little change to the optimized solutions. We initialized the algorithm with a Gaussian distribution with mean 0.1 and a diagonal covariance matrix with variance 0.09 in all dimensions. We ran our parallelized code on two Intel Xeon CPU E5-4640 processors. The optimization for a single movement took about 6 hours of computation time. In Al Borno et al. [2], we show that this method can synthesize movements that replicate kinematic features reported in motor control studies and in experimental three-dimensional reaching data.

### 2.3 Computing Synergies

We incorporated time-invariant muscle synergies in our computational model. According to the time-invariant formulation, the muscle excitations **e** can be decomposed as a linear combinations of synergies **w**_*i*_:

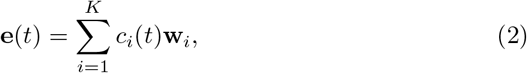

where *c_i_* denote the combination coefficients, *t* denotes time and *K* denotes the number of synergies. When performing trajectory optimization (Sec. 2.2) with synergies, the free variables were the combination coefficients *c_i_* instead of the muscle excitations.

We computed muscle synergies when reaching to a target on a horizontal plane from a chosen initial starting pose (see Fig. 1). The plane was placed approximately 20 cm below the top of the clavicle. We randomly chose a target position on the plane, within a squared area 65 cm on a side and solved a trajectory optimization problem to steer the hand as close as possible to the target. We empirically found that fifteen trajectory optimizations with random targets was sufficient to cover the workspace (i.e., the domain of possible targets). The muscle excitations were stored in a matrix **E** = [**e**_1_,…,**e**_*L*_], where **L** denotes the total number of timesteps from all the synthesized trajectories. We performed dimensionality reduction on **E** with principal component analysis (PCA) to compute the muscle synergies. It has been argued that non-negative matrix factorization (NNMF) computes more physiologically realistic muscle synergies than PCA because it does not allow negative components [52]. In this work, we were mainly interested in studying the computational implications of motor modularity, which could be implemented in different and currently unknown ways. For this reason, we used PCA to compute the modules in our experiments. We used NNMF to assess the sensitivity of our results to the dimensionality reduction technique.

**Figure 1:**
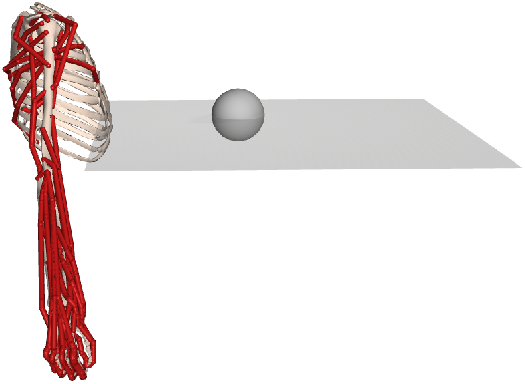
Reaching for a target on a horizontal plane. We solved trajectory optimization problems to reach random targets (gray sphere) on a horizontal plane.

### 2.4 Evaluating Synergies

We next solved trajectory optimization problems to reach new random targets on the plane, except that we constrained the excitations to be linear combinations of synergies by setting the free variables in the optimization to be the synergy combination coefficients, rather than individual muscle excitations. We evaluated how well a task was achieved based on the final value of the cost function in Eq. 1. In addition to the overall cost value, we evaluated the performance of the full and low dimensional controllers in reaching the target (i.e., the Euclidean distance between the final hand position and the target) and minimizing muscle effort (i.e., the sum of muscle activations squared). We chose every third number between 5 and 20 as the number of synergies to study how performance varies with the amount of dimensionality reduction.

We also investigated the potential consequences of muscle synergies for motor skill acquisition by analyzing how muscle synergies impacted the time required to achieve a desired performance level with trajectory optimization. Thus we assumed that optimization time is correlated with motor skill acquisition difficulty (i.e, how much practice is required to achieve a given task). In particular, we compared the number of samples required to achieve the desired performance because sample evaluation was the most computationally expensive operation in the optimization. We said that a desired performance had been achieved when the value of the cost function for the current optimization reached a threshold value, which we set to 0.1. At each iteration in the trajectory optimization,we used the same number of samples as the control dimension (e.g., with 8 synergies, we had 8 samples). We tried using fewer samples per iteration for the full-dimensional optimization (specifically 20 instead of 47), but the optimization failed to converge to the desired motor performance.

We then performed computational simulations to determine whether the synergies computed in the reaching task above generalized to different tasks with performance comparable to the full-dimensional optimization. First, we varied the height of the plane by placing it 60 cm higher and 30 cm lower than its initial position (see Fig. 2). We also tested four different initial upper extremity poses chosen to assess the spectrum from which synergies can generalize (see Fig. 2). We next increased the movement speed by making the motion 1.5 times faster. Finally we tested adding a light (1.75 kg) and heavy (8.7 kg) weight to the hand to model the effect of holding a load. Note that we did not recompute the synergies, as these were fixed from the simulation with the initial plane position and model pose (Sec. 2.3).

**Figure 2:**
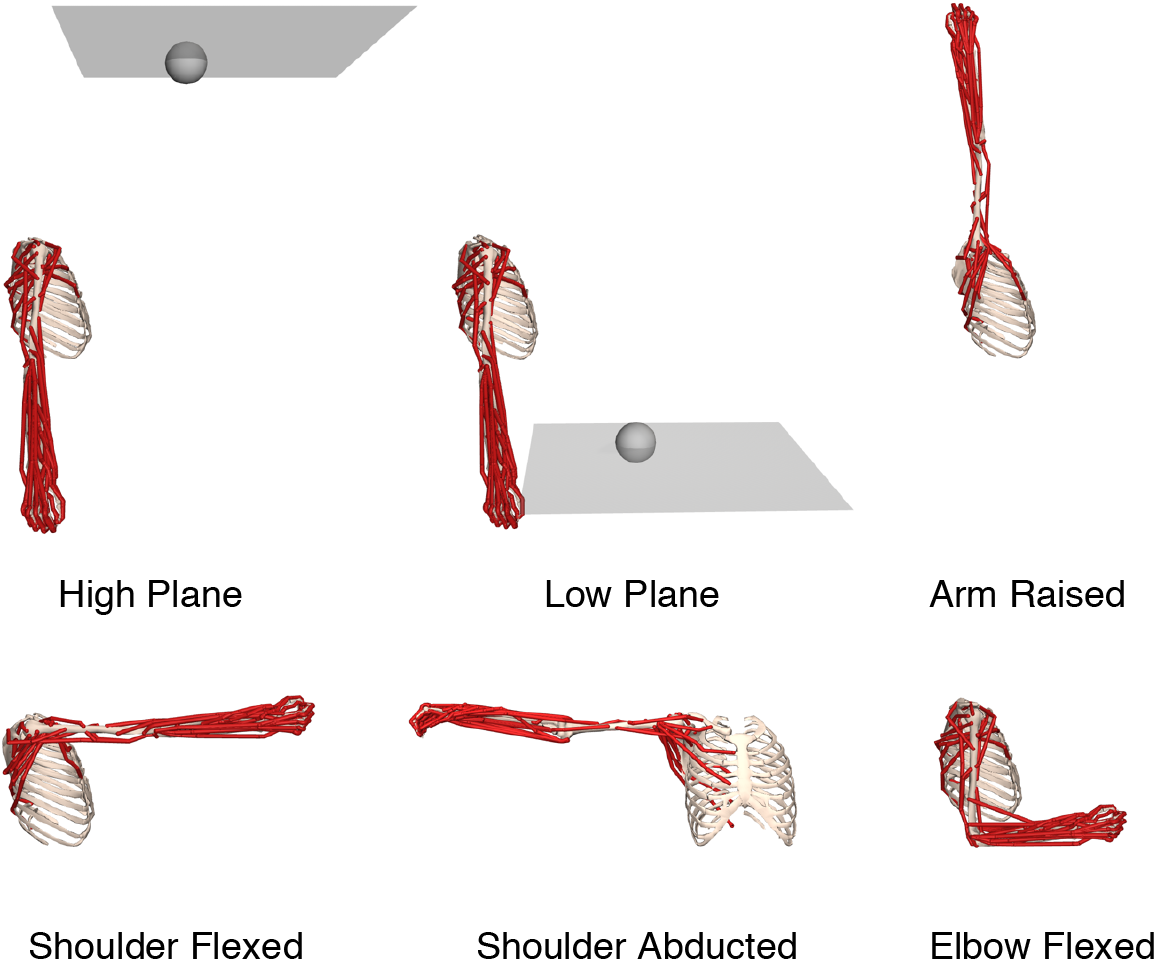
Varying target planes and initial model positions to test the generalizability of synergy-based controllers. We tested the controllers in tasks with a high plane and low plane, with the same initial pose as in the nominal task. We also tested the controllers with varying initial poses, including with the arm raised, shoulder flexed, shoulder abducted, and elbow flexed. For the varying initial poses the targets were in the same plane as in the nominal pose.

### 2.5 Combining Synergies and Independent Muscle Control

Finally, we evaluated a hybrid controller that combined synergies with independent control of each muscle in the model. We compared a model with a hybrid control architecture (i.e., 12 synergies and independent muscle control for a total of 59 control variables) to independent muscle control (i.e., the full-dimensional system with 47 variables) and a model with 12 synergies (and thus 12 control variables). We chose to have 12 synergies in our hybrid architecture to balance between flexibility and learning speed. We used the same number of synergies in the hybrid architecture as in the synergies architecture to aid the comparison. The optimization procedure followed the descriptions in Sec. 2.2 and Sec. 2.3. We set the initial variance in the optimization to 0.0025 for the independent muscles and to 0.09 for the synergies. This means that the optimization prioritized finding a control solution with muscle synergies and started exploring using individual muscles as a secondary step to improve control.

The advantage of the hybrid architecture can be observed when optimizing for a movement outside the space where the synergies are computed. To illustrate this case, we took the initial position of the arm to be as in the experiment with the arm raised (see Fig. 2), while the synergies were computed with the arm in a neutral position (see Fig. 1). To determine the effect of the hybrid architecture on convergence rates (and thus the potential effects on skill acquisition), we also investigated the number of samples required to achieve a desired motor performance (i.e, a cost function value less than 0.1) for the task of reaching to targets in the same plane on which the synergies were created.

## 3 Results

### 3.1 Synergies Do Not Degrade Task Performance

Twenty synergies were required to account for 88% of the variance in 47 muscle excitation patterns for the reaching task in a single horizontal plane (Fig. 3A). While 5 synergies explained only 49% of the variance, with 5 or more synergies, the low-dimensional optimization achieved a performance within 0.011 or 10% of the full-dimensional optimization for new, random reaching targets in the same plane for which synergies were extracted (Fig. 3B). Differences in the overall cost function values on this order represented solutions that are not easily visually distinguishable (see Supplementary Video 1) and are near the variation from run-to-run due to the stochastic nature of the optimization. In particular, over ten runs reaching to the same target, there was variability on the order of 5% of the total cost, 0.5 cm in the final hand position, and of 10% in the movement effort (defined as the sum of muscle activations squared).

**Figure 3:**
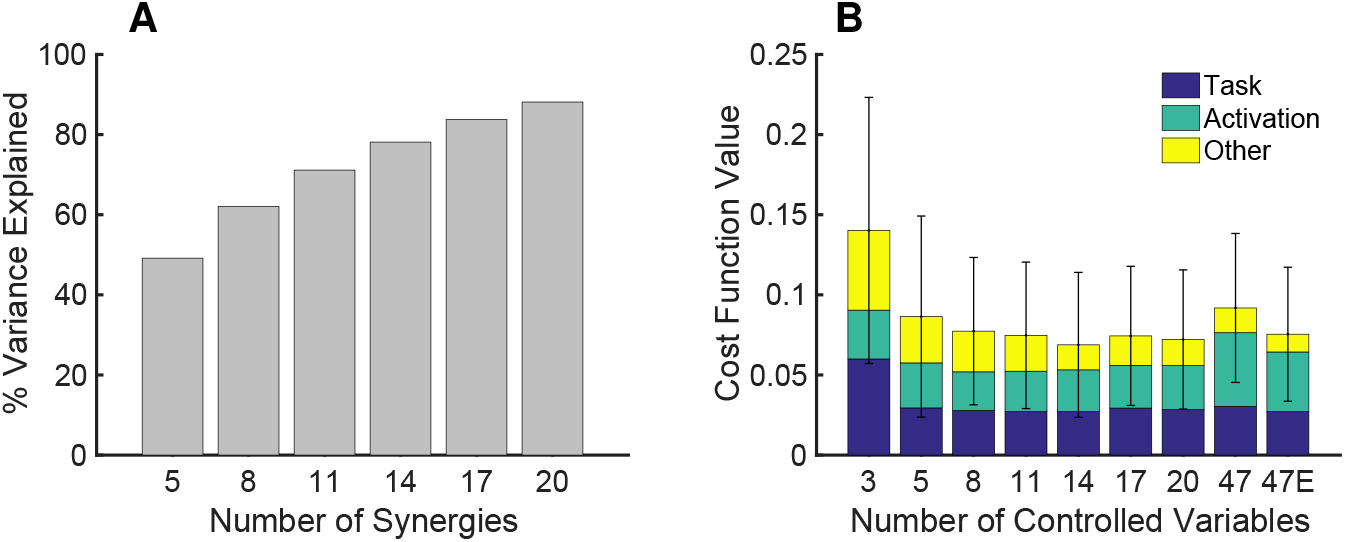
Synergies and task space performance. **A.** Variance in muscle excitations explained versus the number of computed synergies. **B.** Average cost (Eq. 1) for ten random targets in the plane of Fig. 1 when the trajectory optimization was performed with increasing control dimensionality. The cost includes weighted terms to represent task performance (blue), effort (green), and other criteria such as final hand velocity near 0 and joint limit violations. Error bars represent one standard deviation of the cost function value. We set the same population size and the number of iterations in the optimization for the different controllers, except for 47E where we conducted a more exhaustive search by doubling the population size and the number of iterations.

To compare the task performance of the synergy-based controllers to the full-dimensional controller, we set the population size and the number of iterations in CMA-ES to 100 for the synergies and we doubled the population size and the number of iterations for the full-dimensional controller (label “47E” in Fig. 3B). On average, the final hand position with 5 and 8 synergies was 0.6 (1.0) cm and 0.2 (0.8) cm further from the target, respectively, than the hand position with the full-dimensional solution (we provide the standard deviation in parenthesis throughout the paper). While the average error was low for the 5-synergy solution, there were instances where the 5-synergy controller performed poorly. For example, the target error for the worst run with 5 synergies was 2.8 cm further away from the target than the full-dimensional solution. With 3 synergies, the low-dimensional optimization was on average 4.2 (4.7) cm further away from the target than the full-dimensional optimization, an order of magnitude worse than the 5 and 8 synergy solutions. The full-dimensional solution required on average 1.6 (0.3) and 1.3 (0.2) times more effort than the 5 and 8 synergies solutions, respectively. In fact, the low-dimensional optimizations often achieved a lower cost function value since the full-dimensional optimization was more prone to fall in local minima. We found that the optimization tended to converge to movements requiring more effort when more control variables were allowed. This local minima explains why the cost can increase with the number of control variables. When comparing the kinematics of the movements, qualitatively, the synergies produced similar, but simplified curves compared to the full-dimensional system (Fig. 4). We show the movements being performed by the different controllers in Supplementary Video 2.

**Figure 4:**
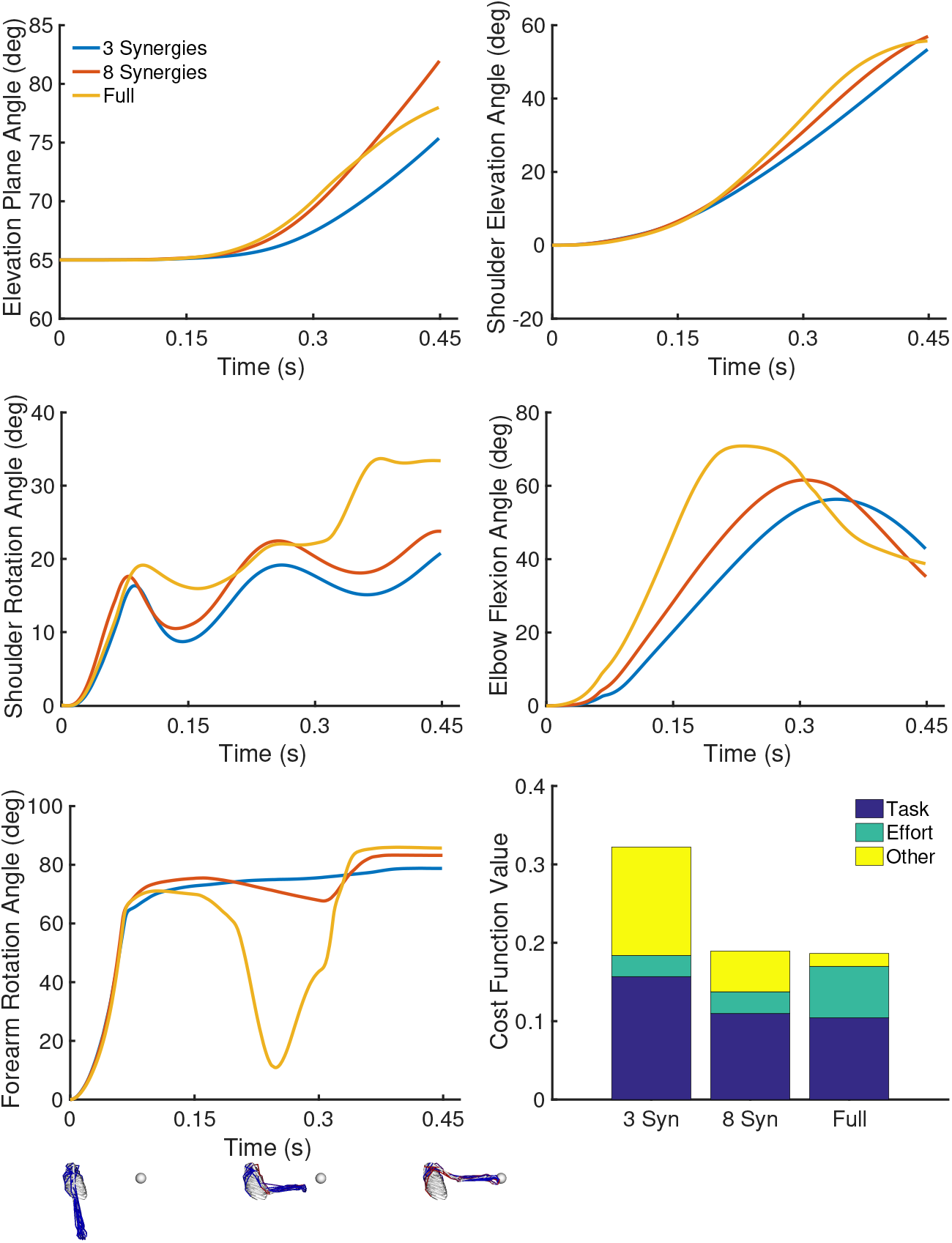
Example kinematics when optimizing reaching to a target with 3 and 8 synergies, and the full-dimensional system. Kinematics are shown for the model’s five degrees-of-freedom, which include the elevation plane, shoulder elevation (or thoracohumeral angle), shoulder rotation (where internal rotation is positive), elbow flexion, and forearm rotation (where pronation is positive) [44]. Kinematics are from a representative run, but the multiple runs were qualitatively similar. We also show some key poses in the movement for the full-dimensional controller. In the bottom rightmost figure, we compare the performance on the cost for each controller for the kinematics in the plots above. We show the proportion of the cost due to the task term, the effort term and all other terms (final velocity near 0, joint limits, etc.; see Eq. 1).

We also conducted a search with the full-dimensional controller using the same population size and number of iterations (100) as for the synergy-based controllers (label “47” in Fig. 3B). This less exhaustive search did not have a large effect on performance. The cost function for the exhaustive search was 0.016 (0.01) lower. On average, this caused the final hand position to be 0.6 (0.4) cm closer to the target, while requiring 16 (20) % less effort than the less exhaustive optimization. The more exhaustive optimization required 25 hours of computation on an Intel Xeon machine with 20 cores for a single movement (as opposed to 6 hours) and yielded relatively small improvements. This indicates that the parameters for the full-dimensional optimization were adequate to converge to a good local minima.

### 3.2 Synergies Facilitate Motor Learning

A controller with 8 or 20 synergies required about 20 and 14 times fewer samples, respectively, to converge to a cost value of 0.1 or less, compared to the optimization with 47 muscles (Fig. 5A). Note that even computing synergies without dimensionality reduction (i.e., having 47 synergies) helped motor learning as the control variables partially encoded the solution, requiring approximately 3 times fewer samples. The 8 synergies optimization required about 18 iterations to achieve the same motor performance (evaluated as the current optimization minimum) that the full-dimensional optimization achieved at about 50 iterations, while requiring fewer samples at each iteration (Fig. 5B). Furthermore, the full-dimensional optimization converged to a worse local minimum than the optimization with 8 synergies. The low-dimensional optimization achieved a final hand position that was 0.21 cm closer to the target and was 2.4 times less costly in terms of the sum of muscle activations squared. We also tested whether randomly generated synergies would speed up learning. None of the five randomly generated, low-dimensional controllers we tested converged to the desired performance (see Fig. S2). This indicates that use of a low dimensional basis alone does not speed up learning.

**Figure 5:**
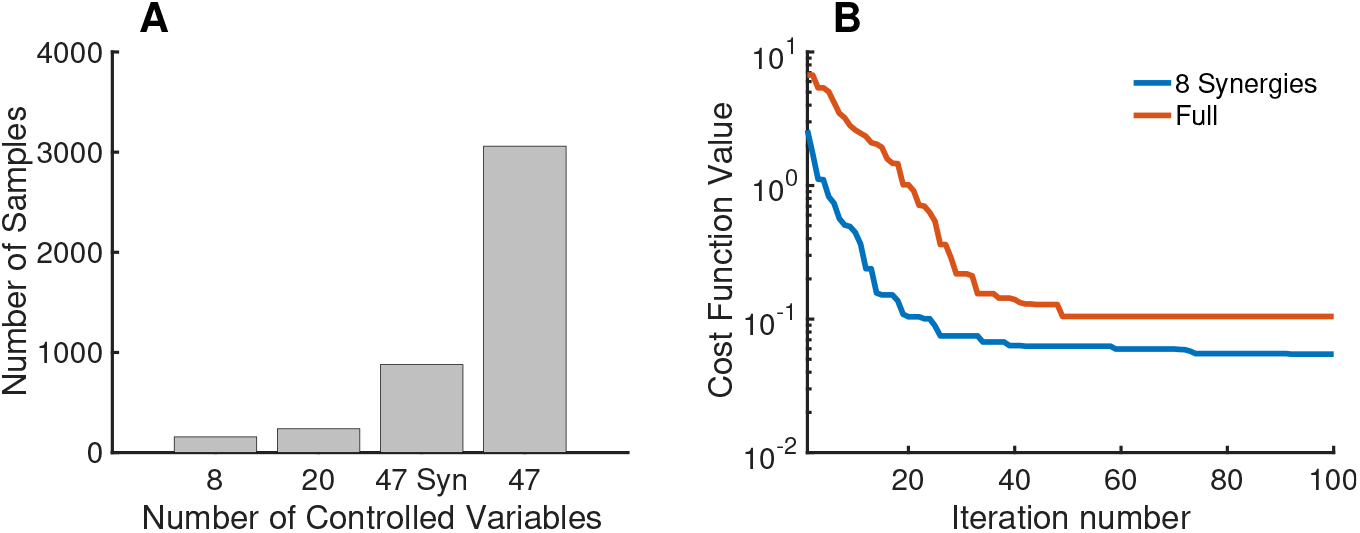
Effect of synergies on search complexity. **A.** The number of samples required to achieve a cost function value less than 0.1 based on the number of controlled variables, averaged over ten random runs. We use “47 Syn” to indicate controlling 47 synergies, instead of controlling the 47 muscles independently. **B.** Convergence rate for the controller with 8 synergies (blue) compared to the full-dimensional system (red).

### 3.3 Synergies Generalize to New Tasks

The optimizations with 8 or more muscle synergies converged to solutions that were similar in cost function value (i.e., on average within 0.005 or better) to the full-dimensional optimization for many, although not all, of the new tasks that we simulated (Fig. 6). When the plane of targets was raised 60 cm higher, the final hand position with 8 synergies was within 1.4 (1.9) cm of the full-dimensional optimization and had 23 (15)% times less effort (Fig. 6A). When the plane was lowered 30 cm and for three of the alternate initial poses (elbow flexed, shoulder flexed and shoulder abducted), all the low-dimensional controllers (i.e., 5 or greater synergies) also achieved an average performance within 0.005 (or better) of the full-dimensional controller, which corresponds to an average difference of 0.36 (0.79) cm in the final hand positions and of 28 (13) % percentage in the total effort (Fig. 6A). While the differences in the percentage of the total effort were large, the effort required for these movements were relatively small;hence, the effort has a small contribution to the overall cost function value. For example, when reaching to the lower plane, the effort terms accounted on average for only 20% of the overall cost. For the initial pose with the arm raised (180 degrees shoulder flexion), the controller needed at least 17 synergies to achieve a comparable (or better) performance to the full-dimensional controller (Fig. 6B). Note that the initial pose in for this task was quite different than the initial pose where the synergies were computed (see Fig. 1). The model was able to execute the reaching task at higher speed and with the 1.75 kg weight using low-dimensional controllers with 8 or more synergies (Fig. 6C). The low-dimensional solution with 8 synergies was on average 0.25 (0.9) cm closer to the target and requires 33 (9) % less total effort. The reaching task with the 8.7 kg weight was a case where the task-specific synergies did not generalize effectively (Fig. 6D), as the full-dimensional solution performed substantially better on Eq. 1. In this case, the full-dimensional optimization was on average 13.0 (7.7) cm closer to the target than the solution with 8 synergies. This is due to the fact that the synergies did not produce sufficient joint torques to hold and move the heavy object because the database of movements from which the synergies were computed did not require large joint torques. We again verified that our results were robust to more exhaustive optimizations (i.e., doubling the population size and the number of iterations in Sec. 2.2 yielded the same trend for the new tasks).

**Figure 6:**
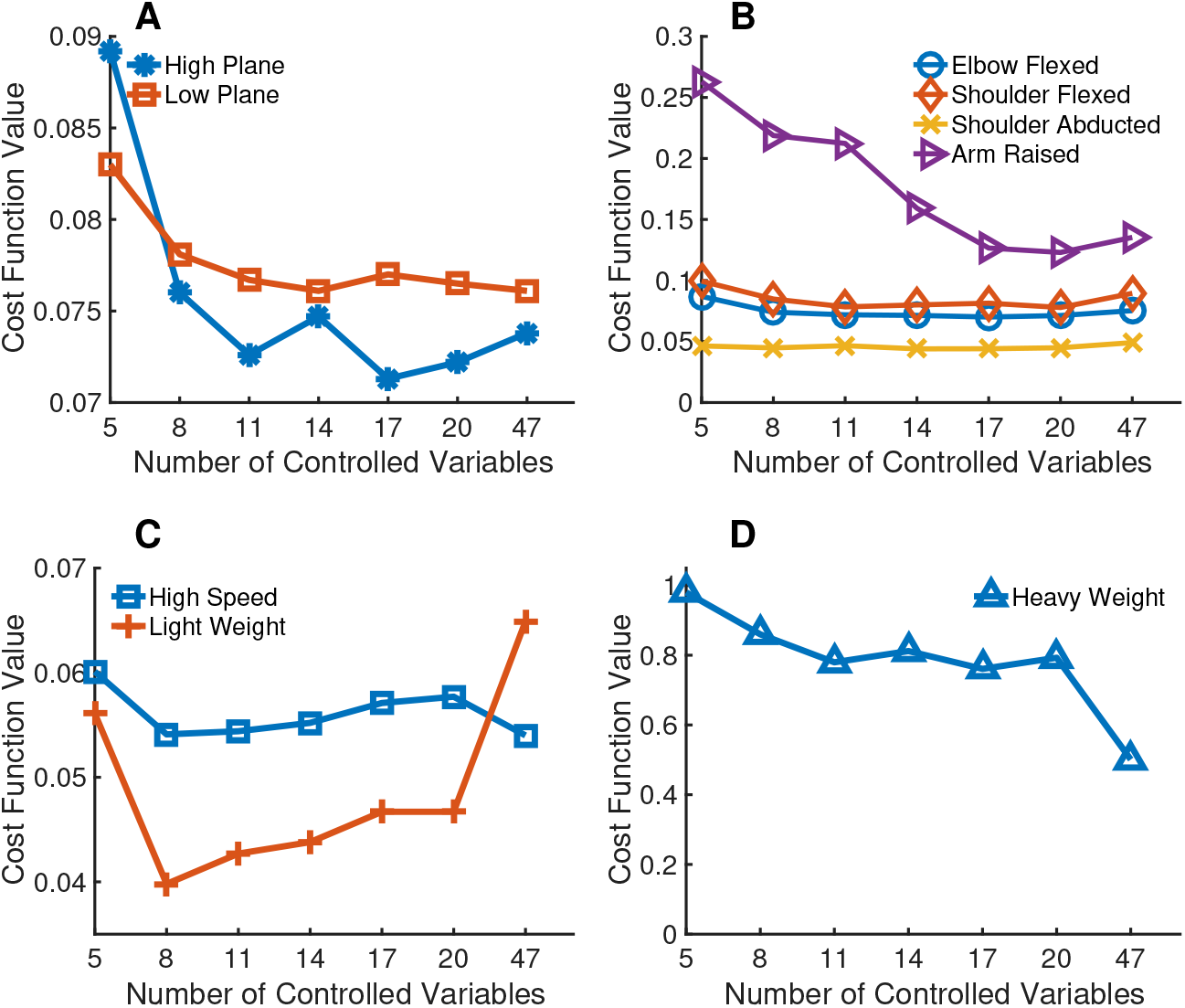
Performance of synergy-based controllers for new tasks. We show the average final cost depending on the number of controlled synergies for several new tasks. **A.** We varied the height of the horizontal plane. The horizontal plane was 60 cm higher and 30 cm lower in the high plane and low plane conditions, respectively. **B.** We varied the initial arm position and the goal was to reach a target on the same horizontal plane as in the nominal task. **C.** The movement was performed 1.5 times faster and we placed a load of 1.75 kg on the hand. **D.** We placed a load of 8.7 kg on the hand. All of these results are the average of ten trajectory optimizations with random targets.

### 3.4 Synergies and Independent Muscle Control

The synergy-only architecture had relatively low performance on the task that required reaching from an arm raised position; however, a hybrid architecture with 12 synergies, along with independent control of the model’s 47 muscles achieved a lower cost than the the full-dimensional optimization, while still being nearly as sample-efficient as the synergy solution. On average, the 12 synergies solution was 6.9 (2.5) cm further away from the target than the full-dimensional solution, while the hybrid solution was 1.8 (2.5) cm further away (see Fig. 7A). On average over ten runs, the hybrid architecture achieved the desired motor performance for the same-plane reaching task using 8.5 fewer samples than the full-dimensional optimization, while the synergies optimization used 20 times fewer samples (see Fig. 7B).

**Figure 7:**
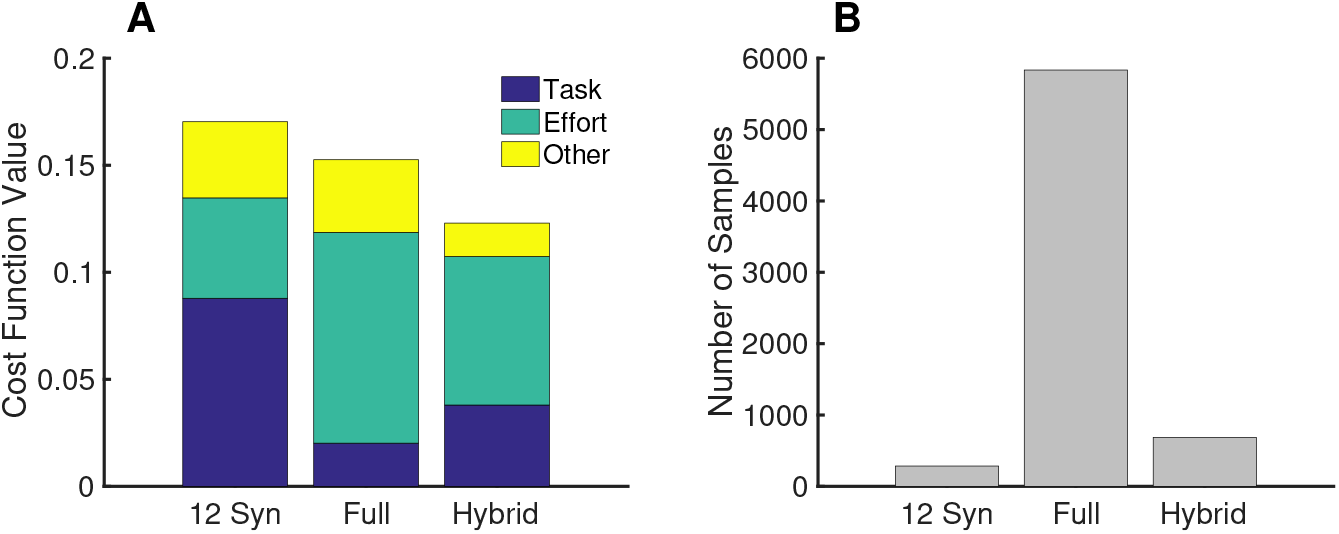
Performance of the hybrid controller. **A.** Average cost function value for 12-synergy, hybrid and full-dimensional architectures across five random targets with the arm raised initial pose (see Fig. 2C). We show the proportion of the total cost due to the task term (blue), the effort term (green), and all other terms (yellow) in Eq. 1. **B.** Number of samples required to achieve a cost function value less than 0.1 based on the number of controlled variables, averaged over ten random runs. Although the hybrid control architecture had 59 variables (12 synergies and 47 independent muscles), it still had a more than eight-fold reduction in the number of samples required to achieve a desired performance when reaching to a target on a horizontal plane (see Fig. 1).

## 4 Discussion

We developed a computational model of upper extremity movement to study the muscle synergy hypothesis for voluntary movement generation. The upper extremity model is more biomechanically realistic than previous work, but it is also more complex to control due to its many muscles and degrees-of-freedom. Our computational experiments indicate that a low-dimensional controller based on muscle synergies does not lead to a significant loss of performance, generates a wide variety of behaviors, and supports the acquisition of skills at a faster rate than a full-dimensional controller (up to a twenty-fold increase in some cases).

A study indicating the converse would pose a serious challenge for the synergy hypothesis. Our results do not imply the existence of synergies, but highlight and quantify their potential benefits. Further speed-ups in motor learning could potentially be achieved with the time-variant formulation of muscle synergies by capturing the time dependence of muscle activations [12], but one would also need to study its effects on movement generalization. We also showed that muscle synergies can be combined with the control of individual muscles to maintain the ability to quickly acquire motor skills, while being able to generalize to more novel tasks.

For our nominal reaching task, 20 synergies were needed to explain 88% of the variance in the muscle excitations. In contrast, experiments with rhesus monkeys indicate that 10 synergies are sufficient to capture 95% of the variance in the reaching EMG data [41]. In human reaching movements on a plane, 4-5 time-varying synergies capture 73-82% of the variance observed in experiments [13]; a larger number of time-invariant synergies is expected to achieve the same variance explained. In experimental studies, EMG is typically only recorded from at most 19 muscles of the upper-extremity due to measurement difficulties. We thus expect a larger number of synergies would be needed to explain most of the variance with our simulations that used 47 muscles to control reaching.Inouyue et al. [28] argue that muscle synergies can cause a drastic reduction in task space performance, and thus percent variance explained may not be a good metric of synergy performance. However, this prior analysis was performed with a simplified biomechanical model, and the authors acknowledge that “having many more muscles will naturally allow the implementation of synergies without such a drastic reduction”. This is the conclusion we have reached in our work. We did not see a significant performance degradation when using 8 synergies that only explained 62% of the variance. Another difference that could explain the divergent results is that the synergies in Inouyue et al. were hand-tuned (and possibly less effective), while our synergies were obtained through optimization. We have also shown that our synergy-based controller often achieved better performance than the full-dimensional optimization, which was more prone to fall in local minima.

The muscle synergies hypothesis is appealing in part because it potentially simplifies the control and learning of movements [6]. Loeb et al. [35] hypothesized that a large action space could facilitate skill acquisition as it would provide an abundance of ways of producing the desired behavior. Our computational study does not support this hypothesis as the large action space made learning more costly by a numerical optimizer. This result is consistent with the experiments by Berger et al. [4] that indicate that it is more difficult (i.e., requires more trials and time) to learn movements that are incompatible with existing experimentally-determined muscle synergies than to learn movements that are consistent with measured synergies. Hagio et al. [24] previously showed that modularity speeds up motor adaptation to rotational perturbations in an isometric force production task. We extend their result by studying reaching movements and by studying the effects of modularity on motor learning, not just adaptation.

Human motor learning can be framed in terms of optimizing a movement policy or reinforcement learning [25]. Our evolutionary algorithm for trajectory optimization (Sec. 2.2) shares similarities with reinforcement learning and can be seen as a scalable alternative [43]. Hence, the number of samples required to achieve a motor performance in our model may provide a first order assessment of the motor learning difficulty of the task in humans. If future research shows that humans learn in ways fundamentally different from our current reinforcement learning models, then our assessment will need to be re-evaluated. State-of-the-art deep reinforcement learning approaches in motor control currently require a prohibitive number of samples to learn new movements, with limited fidelity [34]. It would be interesting to investigate whether modular architectures can help scale these approaches from torque-driven to muscle-based models to achieve greater motion realism. Synergy-based control architectures may also offer an alternative to recent work in computer graphics that attempt to simplify the computational problem by learning realistic torque-driven models instead of muscle-based models directly [29].

There is growing evidence that synergies can generalize across tasks [11, 22, 42, 51], and our study provides further support. We go beyond showing the ability of synergies to reconstruct EMG across tasks and demonstrate that synergies from one reaching task enable task performance in other reaching tasks with accuracy equivalent to a full-dimensional controller. Sohn and colleagues [47] present evidence that generalizability does not arise directly from musculoskeletal or optimality constraints and causes an increase in effort. We complement this study by showing that synergies are generalizable for upper extremity reaching tasks, although we did not see large cost increases for our synergy solutions, likely because we included an effort term for all of our optimizations. The generalization tests designed in our study limit basis generation to a subset of the workspace (i.e., reaching to random targets on a horizontal plane). In an organism, the optimal basis (as might be discovered on evolutionary timescales) can likely only be well identified from a richer dataset, involving a larger workspace (i.e., given the common tasks that an organism must perform). Nevertheless, our study suggests that synergies generalize well outside the workspace, which is valuable independently of how the workspace is defined.

We determined the muscle synergies with PCA as we were interested in analyzing the computational implications of motor modularity without regard to the actual physiological implementation (e.g., in the spinal circuitry [14] or in cortex [21]), which is unknown. We observed a decrease in performance when performing the experiments with non-negative matrix factorization (NNMF), using the solver provided in Kim et al. [32]. For instance, the average cost over five runs when reaching in the horizontal plane was 0.04 higher with NNMF. In task space, this difference amounts to the hand being 2.6 cm further from the target with twice the effort (as measured by the sum of muscle activations squared) compared to the PCA solution. When reaching on the 60 cm higher horizontal plane (outside the space where the synergies were computed), the NNMF cost is 0.069 higher. This reduced performance with NNMF could be due to the fact that PCA generalizes better to data outside the subspace where dimensionality reduction occurs [49]. The synergies determined from NNMF are included in the supplemental material. Similarly to the results reported in d’Avella et al. [11, 13, 45], the first synergy is related to elbow flexion, while synergies 2-4 are related to elbow extension and shoulder flexion.

There are several other modeling simplifications and assumptions that may have impacted our study. First, we did not model spinal circuitry feedback or individual motor units. Second, choosing a different cost function or weights in our cost function might lead to different solutions. However, we chose a cost function and optimization strategy that we have previously shown produces realistic movements with features reported in motor studies [2]. Finally, since our trajectory optimizer is stochastic, it is more likely to return a local minima than the global minimum. In our experiments, the performance of the full-dimensional optimization was not always optimal (e.g., see Fig. 6). The reason is that it is computationally difficult to find the optimal solution with such a high-dimensional system. We chose to terminate our optimizations when the maximum number of iterations was reached for all cases except the learning experiments (where we stopped iterating when a specified performance threshold was achieved). We tested stricter convergence criteria and did not observe markedly different solutions. Our results in Sec. 2.4 indicated that, when employing more exhaustive optimizations, most of the improvements were the in effort term (up to a 16 (20) % improvement in some cases). The standard deviation in the effort term was large, but this is partly due to the fact that this term can have a relatively small contribution to the overall cost function (the optimization aims to reduce the overall cost function value). The improvements in the final hand positions were less than one centimeter. These results indicate that our optimization parameters were sufficient to achieve a good local minima.

Our results in Sec. 3.1 indicate that we should expect variability on the order of half a centimeter in the final hand position and of 10% in the movement effort due to the stochastic optimization. This also limits the resolution of our study: it becomes difficult to assess the implications of synergies at this magnitude. However, we have chosen to perform our analysis on tasks and movements (i.e., three-dimensional upper extremity reaching) where this magnitude is typically not significant.

We have shown that modularity speeds motor learning with our evolutionary optimization algorithm. There may exist other (potentially unknown) algorithms where having independent control of each muscle does not significantly impede motor learning. This touches upon the age-old question of whether intelligent behavior arises because of a prior, possibly innate, structure or because of general learning algorithms. Our results indicate that a modular architecture asa prior structure for reaching movements could improve motor learning, without significantly degrading performance and generalizability. Synergies may be favored by Darwinian natural selection to speed up both motor development and skill acquisition, a hypothesis that is consistent with results in the lower limb that suggest that synergies are conserved across individuals with significantly different motor experiences and even across species [19, 27, 53]. How this prior structure—if it exists—is implemented remains an open question.

## Data accessibility

The code for the computational simulations and our data are available at https://simtk.org/projects/ue-synergies.

## Authors’ contributions

MA conceived and implemented the study, and drafted the manuscript. JH helped interpret results and drafted the manuscript. SD provided edits to the manuscript and supervised all aspects of this work.

## References

[1] Al Borno, M., De Lasa, M., and Hertzmann, A. Trajectory optimization for full-body movements with complex contacts. IEEE transactions on visualization and computer graphics 19, 8 (2013), 1405–1414.

[2] Al Borno, M., Vyas, S., Shenoy, K. V., and Delp, S. L. High-fidelity musculoskeletal modeling reveals a motor planning contribution to the speed-accuracy tradeoff. bioRxiv (2019), 804088.

[3] Alessandro, C., Delis, I., Nori, F., Panzeri, S., and Berret, B. Muscle synergies in neuroscience and robotics: from input-space to task-space perspectives. Frontiers in computational neuroscience 7 (2013).

[4] Berger, D. J., Gentner, R., Edmunds, T., Pai, D. K., and d’Avella, A. Differences in adaptation rates after virtual surgeries provide direct evidence for modularity. Journal of Neuroscience 33, 30 (2013), 12384–12394.

[5] Berniker, M., Jarc, A., Bizzi, E., and Tresch, M. C. Simplified and effective motor control based on muscle synergies to exploit musculoskeletal dynamics. Proceedings of the National Academy of Sciences 106, 18 (2009), 7601–7606.

[6] Bernstein, N. The Co-ordination and Regulation of Movements. Pergamon, 1967.

[7] Bizzi, E., and Cheung, V. C. The neural origin of muscle synergies. Frontiers in computational neuroscience 7 (2013).

[8] Chvatal, S. A., and Ting, L. H. Common muscle synergies for balance and walking. Frontiers in computational neuroscience 7 (2013).

[9] Chvatal, S. A., Torres-Oviedo, G., Safavynia, S. A., and Ting, L. H. Common muscle synergies for control of center of mass and force in nonstepping and stepping postural behaviors. Journal of neurophysiology 106, 2 (2011), 999–1015.

[10] d’Avella, A., and Bizzi, E. Shared and specific muscle synergies in natural motor behaviors. Proceedings of the National Academy of Sciences of the United States of America 102, 8 (2005), 3076–3081.

[11] d’Avella, A., Fernandez, L., Portone, A., and Lacquaniti, F. Modulation of phasic and tonic muscle synergies with reaching direction and speed. Journal of neurophysiology 100, 3 (2008), 1433–1454.

[12] d’Avella, A., and Lacquaniti, F. Control of reaching movements by muscle synergy combinations. Frontiers in computational neuroscience 7 (2013), 42.

[13] d’Avella, A., Portone, A., Fernandez, L., and Lacquaniti, F. Control of fast-reaching movements by muscle synergy combinations. Journal of Neuroscience 26, 30 (2006), 7791–7810.

[14] d’Avella, A., Saltiel, P., and Bizzi, E. Combinations of muscle synergies in the construction of a natural motor behavior. Nature neuroscience 6, 3 (2003), 300–308.

[15] De Groote, F., Jonkers, I., and Duysens, J. Task constraints and minimization of muscle effort result in a small number of muscle synergies during gait. Frontiers in computational neuroscience 8 (2014).

[16] de Rugy, A., Loeb, G. E., and Carroll, T. J. Are muscle synergies useful for neural control? Frontiers in computational neuroscience 7 (2013).

[17] Delp, S. L., Anderson, F. C., Arnold, A. S., Loan, P., Habib, A., John, C. T., Guendelman, E., and Thelen, D. G. Opensim: opensource software to create and analyze dynamic simulations of movement. IEEE transactions on biomedical engineering 54, 11 (2007), 1940–1950.

[18] DeWolf, T., Stewart, T. C., Slotine, J.-J., and Eliasmith, C. A spiking neural model of adaptive arm control. In Proc. R. Soc. B (2016), vol. 283, The Royal Society, p. 20162134.

[19] Dominici, N., Ivanenko, Y. P., Cappellini, G., dAvella, A., Mondi, V., Cicchese, M., Fabiano, A., Silei, T., Di Paolo, A., Giannini, C., ET al. Locomotor primitives in newborn babies and their development. Science 334, 6058 (2011), 997–999.

[20] Flash, T., and Hogan, N. The coordination of arm movements: an experimentally confirmed mathematical model. Journal of neuroscience 5, 7 (1985), 1688–1703.

[21] Gallego, J. A., Perich, M. G., Naufel, S. N., Ethier, C., Solla, S. A., and Miller, L. E. Cortical population activity within a preserved neural manifold underlies multiple motor behaviors. Nature communications 9, 1 (2018), 4233.

[22] Gentner, R., Edmunds, T., Pai, D. K., and d’Avella, A. Robustness of muscle synergies during visuomotor adaptation. Frontiers in computational neuroscience 7 (2013), 120.

[23] Giszter, S. F. Motor primitives—new data and future questions. Current opinion in neurobiology 33 (2015), 156–165.

[24] Hagio, S., and Kouzaki, M. Modularity speeds up motor learning by overcoming mechanical bias in musculoskeletal geometry. Journal of The Royal Society Interface 15, 147 (2018), 20180249.

[25] Haith, A. M., and Krakauer, J. W. Model-based and model-free mechanisms of human motor learning. In Progress in motor control. Springer, 2013, pp. 1–21.

[26] Hansen, N. The cma evolution strategy: a comparing review. Towards a new evolutionary computation (2006), 75–102.

[27] Hart, C. B., and Giszter, S. F. A neural basis for motor primitives in the spinal cord. Journal of Neuroscience 30, 4 (2010), 1322–1336.

[28] Inouye, J. M., and Valero-Cuevas, F. J. Muscle synergies heavily influence the neural control of arm endpoint stiffness and energy consumption. PLoS computational biology 12, 2 (2016), e1004737.

[29] Jiang, Y., Van Wouwe, T., De Groote, F., and Liu, C. K. Synthesis of biologically realistic human motion using joint torque actuation. ACM Transactions on Graphics (TOG) 38, 4 (2019), 1–12.

[30] Kargo, W. J., and Giszter, S. F. Individual premotor drive pulses, not time-varying synergies, are the units of adjustment for limb trajectories constructed in spinal cord. Journal of Neuroscience 28, 10 (2008), 2409–2425.

[31] Kargo, W. J., Ramakrishnan, A., Hart, C. B., Rome, L. C., and Giszter, S. F. A simple experimentally based model using proprioceptive regulation of motor primitives captures adjusted trajectory formation in spinal frogs. Journal of neurophysiology 103, 1 (2010), 573–590.

[32] Kim, J., and Park, H. Fast nonnegative matrix factorization: An activeset-like method and comparisons. SIAM Journal on Scientific Computing 33, 6 (2011), 3261–3281.

[33] Lafreniere-Roula, M., and McCrea, D. A. Deletions of rhythmic motoneuron activity during fictive locomotion and scratch provide clues tothe organization of the mammalian central pattern generator. Journal of neurophysiology 9f, 2 (2005), 1120–1132.

[34] Lillicrap, T. P., Hunt, J. J., Pritzel, A., Heess, N., Erez, T., Tassa, Y., Silver, D., and Wierstra, D. Continuous control with deep reinforcement learning. arXiv preprint arXiv:1509.02971 (2015).

[35] Loeb, G. E. Optimal isnt good enough. Biological cybernetics 106, 11-12 (2012), 757–765.

[36] McGowan, C. P., Neptune, R. R., Clark, D. J., and Kautz, S. A. Modular control of human walking: adaptations to altered mechanical demands. Journal of biomechanics 43, 3 (2010), 412–419.

[37] Meyer, A. J., Eskinazi, I., Jackson, J. N., Rao, A. V., Patten, C., and Fregly, B. J. Muscle synergies facilitate computational prediction of subject-specific walking motions. Frontiers in bioengineering and biotechnology 4 (2016), 77.

[38] Muceli, S., Boye, A. T., d’Avella, A., and Farina, D. Identifying representative synergy matrices for describing muscular activation patterns during multidirectional reaching in the horizontal plane. Journal of neurophysiology 103, 3 (2010), 1532–1542.

[39] Nazarpour, K., Barnard, A., and Jackson, A. Flexible cortical control of task-specific muscle synergies. Journal of Neuroscience 32, 36 (2012), 12349–12360.

[40] Neptune, R. R., Clark, D. J., and Kautz, S. A. Modular control of human walking: a simulation study. Journal of biomechanics 42, 9 (2009), 1282–1287.

[41] Overduin, S. A., d’Avella, A., Roh, J., Carmena, J. M., and Bizzi, E. Representation of muscle synergies in the primate brain. Journal of Neuroscience 35, 37 (2015), 12615–12624.

[42] Roh, J., Rymer, W. Z., and Beer, R. F. Robustness of muscle synergies underlying three-dimensional force generation at the hand in healthy humans. Journal of neurophysiology 107, 8 (2012), 2123–2142.

[43] Salimans, T., Ho, J., Chen, X., Sidor, S., and Sutskever, I. Evolution strategies as a scalable alternative to reinforcement learning. arXiv preprint arXiv:1703.03864 (2017).

[44] Saul, K. R., Hu, X., Goehler, C. M., Vidt, M. E., Daly, M., Velisar, A., and Murray, W. M. Benchmarking of dynamic simulation predictions in two software platforms using an upper limb musculoskeletal model. Computer methods in biomechanics and biomedical engineering 18, 13 (2015), 1445–1458.

[45] Scano, A., Chiavenna, A., Malosio, M., Molinari Tosatti, L., and Molteni, F. Muscle synergies-based characterization and clustering of poststroke patients in reaching movements. Frontiers in bioengineering and biotechnology 5 (2017), 62.

[46] Shadmehr, R. Distinct neural circuits for control of movement vs. holding still. Journal of neurophysiology 117, 4 (2017), 1431–1460.

[47] Sohn, M. H., and Ting, L. H. Suboptimal muscle synergy activation patterns generalize their motor function across postures. Frontiers in computational neuroscience 10 (2016), 7.

[48] Thomas, P. S., and Barto, A. G. Motor primitive discovery. In Development and Learning and Epigenetic Robotics (ICDL), 2012 IEEE International Conference on (2012), IEEE, pp. 1–8.

[49] Ting, L. H., and Chvatal, S. A. Decomposing muscle activity in motor tasks. Motor Control Theories, Experiments and Applications. Oxf. Univ. Press, New York (2010), 102v–138.

[50] Torres-Oviedo, G., Macpherson, J. M., and Ting, L. H. Muscle synergy organization is robust across a variety of postural perturbations. Journal of neurophysiology 96, 3 (2006), 1530–1546.

[51] Torres-Oviedo, G., and Ting, L. H. Subject-specific muscle synergies in human balance control are consistent across different biomechanical contexts. Journal of neurophysiology 103, 6 (2010), 3084–3098.

[52] Tresch, M. C., Cheung, V. C., and d’Avella, A. Matrix factorization algorithms for the identification of muscle synergies: evaluation on simulated and experimental data sets. Journal of neurophysiology 95, 4 (2006), 2199–2212.

[53] Yang, Q., Logan, D., and Giszter, S. F. Motor primitives are determined in early development and are then robustly conserved into adulthood. Proceedings of the National Academy of Sciences 116, 24 (2019), 12025–12034.

[54] Zajac, F. E. Muscle and tendon: properties, models, scaling, and application to biomechanics and motor control. Critical reviews in biomedical engineering 17, 4 (1989), 359–411.

