## Supplemental Material for "The Effects of Motor Modularity on Performance, Learning, and Generalizability in Upper-Extremity Reaching: a Computational Analysis"

| Nbr | Muscle |
|---|---|
| 1 | Deltoid Anterior |
| 2 | Deltoid Medial |
| 3 | Deltoid Posterior |
| 4 | Supraspinatus |
| 5 | Infraspinatus |
| 6 | Subscapularis |
| 7 | Teres minor |
| 8 | Teres major |
| 9 | Pectoralis major Clavicular |
| 10 | Pectoralis major Sternal |
| 11 | Pectoralis major Ribs |
| 12 | Latissimus dorsi Thoracic |
| 13 | Latissimus dorsi Lumbar |
| 14 | Latissimus dorsi Iliac |
| 15 | Coracobrachialis |
| 16 | Elbow Triceps Long |
| 17 | Elbow Triceps Lateral |

| | |
|---|---|
| 18 | Elbow Triceps Medial |
| 19 | Anconeuse |
| 20 | Supinator |
| 21 | Biceps Brachialis |
| 22 | Biceps Brachioradialis |
| 23 | Extensor carpi radialis longus |
| 24 | Extensor carpi radialis brevis |
| 25 | Extensor carpi ulnaris |
| 26 | Flexor carpi radialis |
| 27 | Flexor carpi ulnaris |
| 28 | Palmaris longus |
| 29 | Pronator teres |
| 30 | Pronator quadratus |
| 31 | Flexor digitorum superficialis Digit 5 |
| 32 | Flexor digitorum superficialis Digit 4 |
| 33 | Flexor digitorum superficialis Digit 3 |
| 34 | Flexor digitorum superficialis Digit 2 |
| 35 | Flexor digitorum profundus Digit 5 |
| 36 | Flexor digitorum profundus Digit 4 |
| 37 | Flexor digitorum profundus Digit 3 |
| 38 | Flexor digitorum profundus Digit 2 |
| 39 | Extensor digitorum communis Digit 5 |
| 40 | Extensor digitorum communis Digit 4 |
| 41 | Extensor digitorum communis Digit 3 |
| 42 | Extensor digitorum communis Digit 2 |
| 43 | Extensor digiti minimi |
| 44 | Extensor indicis propius |
| 45 | Extensor pollicis longus |
| 46 | Extensor pollicis brevis |
| 47 | Flexor pollicis longus |

Table S1: **Muscle for each number.**

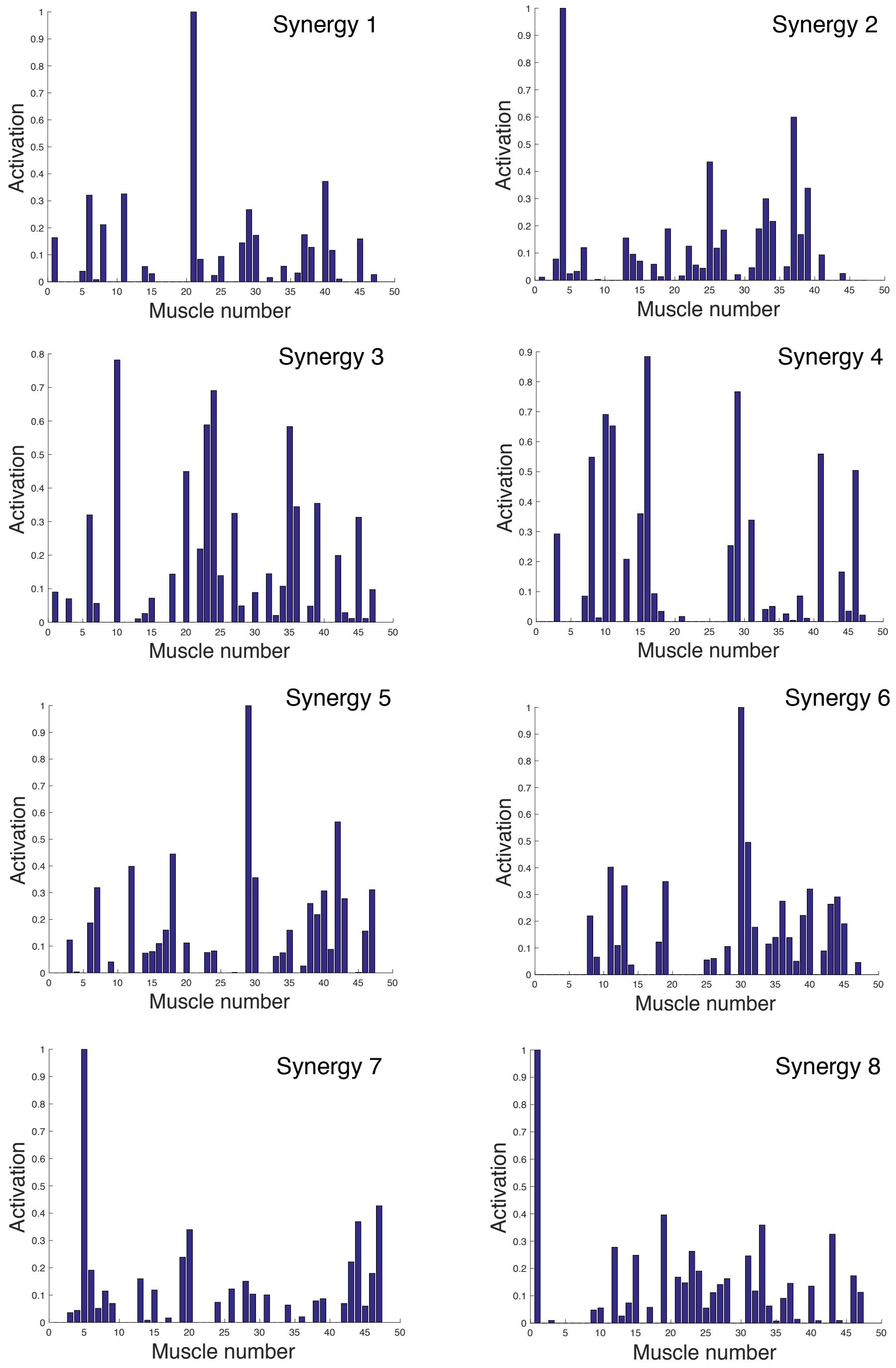


Figure S1: **First eight synergies computed by NNMF..**

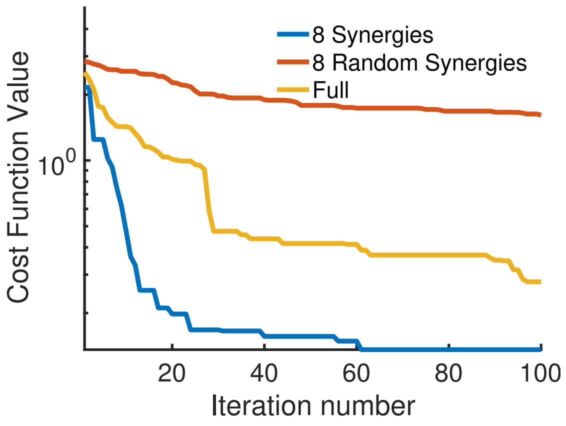


Figure S2: **Effect of synergies on convergence rates.** Convergence rate for the controller with 8 synergies (blue) compared to 8 random synergies (red) and the full-dimensional system (yellow) for a representative run.
